# Sympathetic nerve activity promotes cardiomyocyte cell-cycle arrest and binucleation

**DOI:** 10.1101/124347

**Authors:** Li Chen, Alexander Y. Payumo, Kentaro Hirose, Rachel B. Bigley, Jonathan Lovas, Rejji Kuruvilla, Guo N. Huang

## Abstract

Adult mammalian hearts typically have little capacity to regenerate after injuries such as myocardial infarction. In contrast, neonatal mice during the first week of life possess an incredible ability to regenerate their hearts, though this capacity is lost shortly after birth. The physiological triggers mediating this transition remains poorly understood. In this study, we demonstrate that sympathetic nerve activity promotes cardiomyocyte cell-cycle arrest and binucleation. In mice hearts lacking sympathetic nerve inputs, we observe increased mononucleated cardiomyocyte numbers and elevated cardiomyocyte proliferation. Additionally, increased cardiomyocyte mononucleation and proliferation are observed in mice with genetic and pharmacological inhibition of β-adrenergic receptors (βARs), which mediate sympathetic nerve signaling. Using *in vitro* cultures of neonatal cardiomyocytes, we demonstrate that activation of β-adrenergic receptors results in decreased cardiomyocyte proliferation that is mediated through cyclic AMP-dependent protein kinase (PKA) signaling. Taken together, these results suggest that sympathetic nerve activity may play a role in limiting the ability of mammalian hearts to regenerate by restricting cardiomyocyte proliferation and promoting cytokinesis failure leading to multinucleation.

## INTRODUCTION

Lower vertebrates such as adult newts and zebrafish, and even mammals at embryonic and neonatal stages possess remarkable abilities to robustly regenerate their hearts.^1^ Previous studies have demonstrated vigorous cardiomyocyte proliferation in injured newt hearts,^2–4^ particularly when the hearts were injured at the base of the ventricles.^5^ Lizard cardiomyocytes demonstrate robust proliferation and regeneration after injury.^6^ Adult zebrafish are capable of complete heart regeneration after surgical removal of up to 20% of the ventricle as well as in cryoinjury and genetic cell-ablation models.^7,8^ Recently, an amazing capacity for complete cardiac regeneration has been observed in neonatal mice^9^ and even humans.^10^ However, this capacity is lost shortly after birth in one-week old mice^9^, consistent with the inability of adult humans to regenerate their hearts after myocardial infarction.^1,9,11–13^ In line with this, most mammalian organs and appendages including brain, skin, and limbs have reduced regenerative potential during postnatal growth.^9,14–18^ Understanding the evolutionary and developmental mechanisms regulating regenerative potential could lead to new therapeutic strategies in regenerative medicine; however, our current understanding of these mechanisms is limited.

In adult zebrafish and neonatal mice, heart regeneration occurs primarily through proliferation of pre-existing cardiomyocytes rather than expansion and differentiation of stem cell populations.^1,11,19,20^ Unfortunately, mammalian cardiomyocytes withdraw from the cell-cycle shortly after birth.^1^ In newborn mice, cardiomyocytes undergo a final round of DNA duplication and division during the first weeks of life.^12^ However, due to cytokinesis failure, approximately 90 percent of murine cardiomyocytes binucleate rather than completely divide into individual cells. The increase in heart size from neonatal to adult stages is predominantly due to cardiac hypertrophy with cardiomyocyte renewal rates at less than one percent per year.^1^ In contrast, greater than 95 percent of adult zebrafish and newt cardiomyocytes are mononucleated and retain proliferative activity.^21,22^ Because of this, it is hypothesized that loss of mammalian cardiomyocyte proliferative activity during neonatal stages restricts cardiac regenerative potential in adulthood.

Significant progress has been made in identifying intrinsic regulators of cardiomyocyte proliferation including cyclins, microRNAs, transcription factors and cofactors YAP,^23–28^TBX20^29–34^, and MEIS1^35^. A significant gap in our understanding is in understanding what upstream extrinsic triggers drive permanent mammalian cardiomyocyte cell-cycle arrest. Exposure to postnatal oxygen levels and accompanying increases in oxidative stress has been identified as a driver of cardiomyocyte cell-cycle arrest.^36^ However, only a marginal increase in mononucleated cardiomyocytes of less than five percent was observed in animals chemically or genetically manipulated to reduce oxidative stress, suggesting that other pathways are likely to also regulate this process.

In this study, we analyze the effects of the sympathetic nervous system in cardiomyocyte cell-cycle control. Sympathetic nerves regulate the rate and force of cardiomyocyte contractility through activation of β-adrenergic receptors (βARs) expressed in the innervated heart.^37^ While sympathetic nerve innervation is observed in murine hearts by embryonic stage 13.5 (E13.5),^38^ studies in rats suggest that sympathetic nerve activation does not occur until after birth.^39^ As this coincides with the timing of cardiomyocyte binucleation and cell-cycle arrest,^40^ we hypothesized that sympathetic nerve activity could be an extrinsic factor mediating this transition. Here we show that mice lacking sympathetic nerve inputs retain significantly higher levels of mononucleated cardiomyocytes with increased proliferative potential. Additionally, we demonstrate that genetic and chemical inactivation of βARs similarly result in increased cardiomyocyte mononucleation and proliferation. Finally, *in vitro* experiments in cultured neonatal cardiomyocytes show that pharmacological activation of βARs inhibits cardiomyocyte proliferation through a cyclic AMP-dependent protein kinase (PKA)-dependent mechanism.

## METHODS

### Animals

All protocols have been approved by the Institutional Animal Care and Use Committee of the University of California, San Francisco. *TH-Cre; TrkA^f/f^* mice have been described previously.^41^ βAR TKO mice (beta-less) have also been recently published.^42^ Propranolol (Sigma, 40543) was dissolved at a concentration of 40 mg/mL dissolved in citrate buffer (pH = 3) and then diluted 10-fold with 0.9% saline solution at a working concentration of 4 mg/mL. CRL:CD1 pups (ICR, Charles River) were injected subcutaneously once a day from postnatal stage 0 (P0) to P14 at a dose of 4 μg/g bodyweight propranolol or equal volume saline as control.

### Cardiomyocyte dissociation

Cardiomyocytes from *TH-Cre; TrkA^f/f^* and *TrkA^f/f^* littermate controls were dissociated by fixing harvested hearts for two hours in 3.7% formaldehyde at room temperature. The hearts were then washed for 10 minutes twice in PBS, cut into small pieces and incubated with digestion buffer [1.8 mg/mL Collagenase B (Fisher), 2.4 mg/mL Collagenase D (Fisher), 0.5x Hank’s Buffer (Corning), 1 mM MgCl_2_, and 4 mM CaCl_2_] with rocking agitation at 37°C overnight. The mixture was then collected and centrifuged at 200 x g for five minutes to pellet dissociated cardiomyocytes. The cells were then resuspended in PBS, adhered to Superfrost Plus slides (Fisher), stained with DAPI, and imaged.

Cardiomyocytes dissociated from P14 propranolol- or saline-injected hearts and one-year-old βAR TKO mice or wildtype controls were isolated by Langendorff perfusion as previously described^43^ with slight modifications. After perfusion of hearts for 15 to 20 minutes with 2 mg/mL collagenase II (Worthington) in perfusion buffer [120.4 mM NaCl, 14.7 mM KCl,0.6 mM KH_2_PO_4_, 0.6 Na_2_HPO_4_*7H_2_O, 1.2 mM MgSO_4_*7H_2_O, 10 mM HEPES, 4.6 mM NaHCO_3_, 30 mM Taurine, 5.5 mM glucose, and 10 mM 2,3-butanedione monoxime], partially digested hearts were gently triturated to release individual cardiomyocytes. Dissociated cells were then resuspended in digestion buffer, mixed with an equal volume of 3.7% formaldehyde, and allowed to fix for 10 to 15 minutes. The dissociated cardiomyocytes were then pelleted by centrifugation at 200 × g for five minutes and resuspended in PBS. Cardiomyocytes were then adhered to Superfrost Plus slides, stained with DAPI, and imaged.

### Cryosectioning

Hearts were harvested and blood manually removed through gentle pumping. Extraneous tissues were excised leaving only atria and ventricles intact. Hearts were then briefly soaked in chilled 30% sucrose in PBS and then mounted in cryomolds (TissueTek) containing O.C.T. compound (TissueTek) on ice, allowed to equilibrate for 10 minutes, and then flash-frozen in liquid nitrogen. Tissue blocks were then sectioned to 5-mm thickness using a Leica CM3050 S cryostat, allowed to dry, and then stored at −80°C. Prior to staining, sections were fixed for 10 to 15 min in 3.7% formaldehyde and washed with PBS. After fixation, sections were washed three times in PBS prior to antibody staining.

### Neonatal cardiomyocyte isolation and culture

Our culture protocol for neonatal ventricular cardiomyocytes was adapted from one previously established for adult zebrafish cardiomyocyte culture.^44^ Ventricles were isolated from P1 to P3 neonatal mice and rinsed in perfusion buffer (same as used for cardiomyocyte dissociation) Each heart was then cut into four small pieces to facilitate digestion and transferred into 2-mL Eppendorf tubes (at most two hearts per tube) containing 1 mL digestion buffer (2 mg/mL collagenase II dissolved in perfusion buffer). Tubes were incubated with rocking agitation at 37°C incubator for 30 minutes. Heart pieces were allowed to settle, supernatant discarded, and fresh pre-warmed digestion buffer replaced. Samples were then incubated for 15 minutes at 37°C, supernatants again discarded, and pre-warmed digestion buffer replaced. Next, samples were incubated for 15 minutes and this time, the supernatant was collected and 1/10^th^ volume FBS was added to neutralize the digestion. Fresh pre-warmed digestion buffer was added to the undigested heart pieces and subsequent collections were taken every 10 minutes until complete digestion of the heart was observed. Pooled collections were then centrifuged at 200 × g for five minutes at 4°C. Pelleted cells were then washed in perfusion buffer containing increasing concentrations of CaCl_2_ (12.5 μm, 62 μm, 112 μm, 212 μm, 500 μm, 1000 μm). Finally, cells were resuspended in culture media consisting of high glucose DMEM (UCSF cell culture facility) with 20% FBS (UCSF cell culture facility), 5% horse serum (Corning Cellgro), and primocin (InvivoGen). Cells were then pre-plated in a 6-well culture plate for 1 hour at 37°C with 5% CO_2_ to enrich for cardiomyocytes prior to plating on 0.2% gelatin-coated 96-well plates (Corning Life Sciences). In 100 μL volumes, 20,000 to 25,000 cells were plated per each well of a 96-well plate. Culture media was replaced at least once every two days as needed.

### Chemical treatments

Chemicals were added to cultured cardiomyocytes 24 hours after initial plating at the following concentrations: 5 μM 5-ethynyl-2’-deoxyuridine (SCBT, sc-284628); 10 μM norepinephrine (Sigma, A0937); 10 μM isoproterenol (Sigma, I6504); 100 μM phenylephrine (Sigma, P6126); 10 μM propranolol; 1 μM forskolin (gift from Roshanak Irannejad) and 1 μM KT5720 (EMD-Millipore, 80052-786). After incubation in the compounds for 48 hours, cardiomyocytes were washed briefly with PBS, fixed in 3.7% formaldehyde for 10 to 15 minutes, and washed again with PBS.

### EdU detection

Fixed cultured cardiomyocytes were permeabilized for 20 minutes with PBS containing 0.2% Triton-X 100 (PBST), washed briefly with PBS and blocked in 3% BSA for 20 minutes at room temperature. Cells were then incubated in staining solution [0.1 M ascorbic acid, 1 mM CuSO4, 100 mM Tris pH = 8, and 10 uM sulfocyanine-5 azide dye (Lumiprobe)] for 20 minutes and then washed in PBS.

### Immunohistochemistry

Fixed tissue sections or cultured cells were blocked for at least 1 hour in 10% normal donkey serum (Jackson Immuno Research) in PBST. Primary antibody incubation of 1:200 rabbit anti-PCM1 (SCBT, SC-67204), 1:200 rat anti-Ki67 eFlour 570 (eBiosciences, 41-5698-80), and 1:400 mouse anti-cTnT (Thermo Scientific, MS295P1) was performed at 4 □ overnight. Samples were washed three times with PBST, incubated with fluorophore-conjugated secondary antibodies for at least 2 hours at room temperature, washed three times with PBST, and then stained with DAPI.

### Imaging and image processing

Tissue sections and cultured cells were imaged on an inverted epifluorescent microscope (Nikon Eclipse Ti) or an upright confocal set-up (Leica SPE). Images were processed in FIJI (https://fiji.sc/) or Photoshop (Adobe) and linear adjustments were made to brightness and contrast settings.

### Image quantification

Images were quantified manually with either FIJI using the Cell Counter plug-in or Photoshop. For nucleation analysis, approximately 200 unambiguous cardiomyocytes were analyzed per sample. For tissue sections, Ki67-(+) cardiomyocyte nuclei were quantified in three independent sections per sample and averaged. For cultured cardiomyocytes, an average of 750 cardiomyocyte nuclei were analyzed per condition.

### Statistics

P-values were obtained using the Student’s t Test when comparing two independent samples. For multiple comparisons, one-way analysis of variance (ANOVA) was performed using Prism 7 (GraphPad).

## RESULTS

### Sympathetic nerve ablation increases cardiomyocyte mononucleation and proliferation

To assess the role of sympathetic nerves in the regulation of cardiomyocyte proliferation and nucleation, we utilized a previously established genetic approach to ablate sympathetic nerve innervation.^38^ As the survival of sympathetic neurons requires expression of the nerve growth factor receptor *Tropomyosin receptor kinase A* (*TrkA*),*^45^* we crossed mice harboring a conditional (floxed) *TrkA* allele (*TrkA^f/f^*) with mice containing *Tyrosine hydroxylase* (*TH*)*-driven* Cre recombinase (*TH-Cre*). In *TH-Cre; TrkA^f/f^* mice, sympathetic neurons are specifically depleted of *TrkA*, fail to survive, and are depleted.^38^

By postnatal stage 14 (P14), approximately 90% of murine cardiomyocytes are binucleated and exhibit minimal levels of proliferation while approximately 10% remain as mononucleated cardiomyocytes.^40^ Using this developmental stage as a point of comparison, we harvested hearts from both *TH-Cre; TrkA^f/f^* mice and *TrkA^f/f^* sibling controls and dissociated the tissues into isolated single-cells (Fig. 1A-B). From these dissociated samples, we observed intact cardiomyocytes containing a single nucleus (mononucleated), two nuclei (binucleated), and greater than or equal to three nuclei (polynucleated). Quantification of cardiomyocyte nucleation state under these conditions revealed that *TH-Cre; TrkA^f/f^* mice possessed elevated mononucleated cardiomyocyte numbers (23.0%, n = 3) compared to that observed in *TrkA^f/f^* sibling controls (13.8%, n = 3).

**Figure 1.**
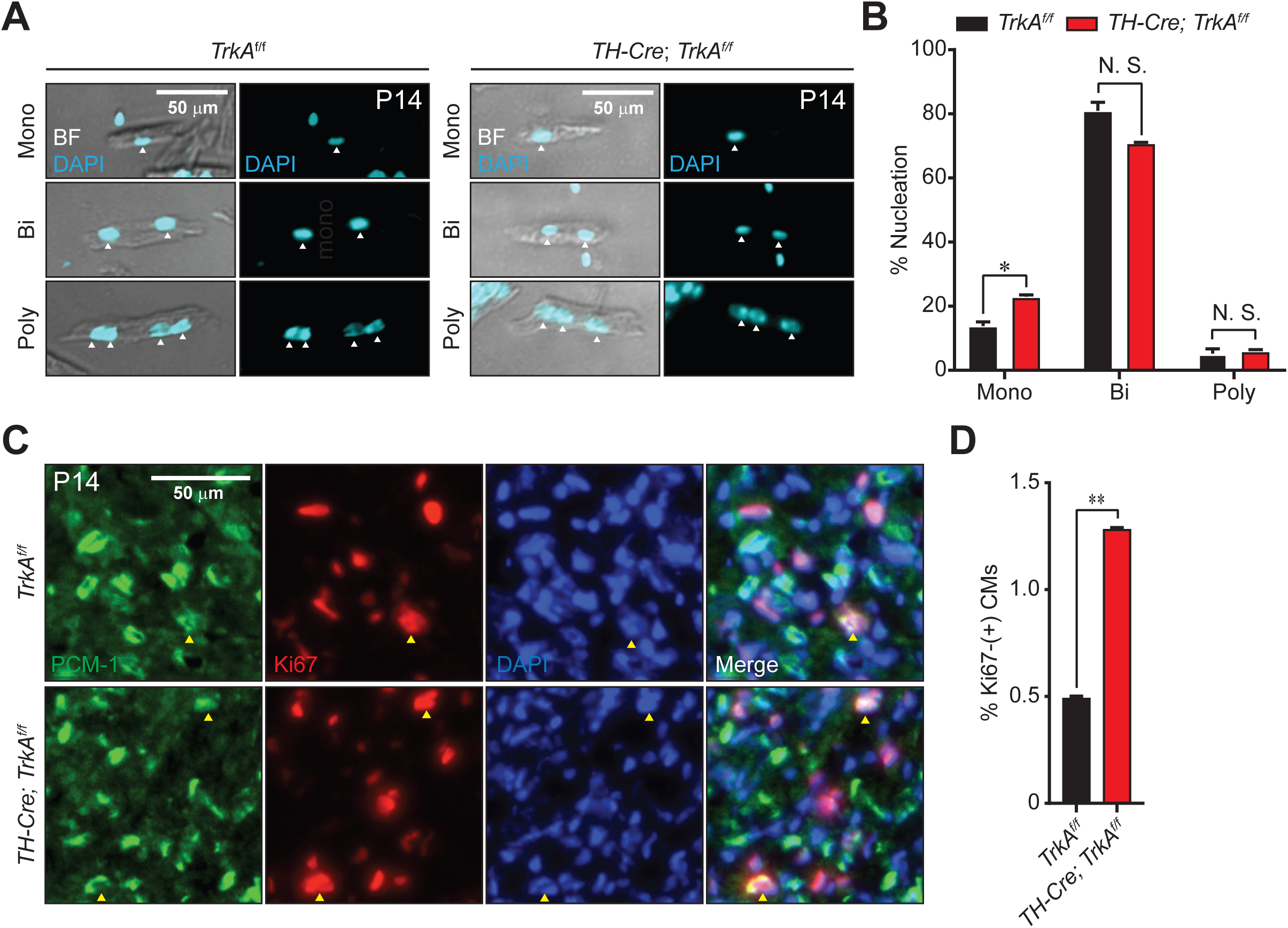
Sympathetic nerve ablation increases cardiomyocyte mononucleation and proliferation *in vivo*.

**A)** Representative micrographs and **B)** quantification of mononucleated (Mono), binucleated (Bi), and polynucleated (Poly) cardiomyocytes dissociated from *TH-Cre; TrkA^f/f^* mice (n = 3) and *TrkA^f/f^* littermate controls (n = 3). White arrowheads indicate cardiomyocyte-specific nuclei. **C)** Representative micrographs and **D)** quantification of 5 μm-thick sections in mutant (n=3) and control (n =3) mice stained with PCM-1 (cardiomyocyte nuclei), Ki67 (proliferative cells), and DAPI (all nuclei). Yellow arrowheads indicate PCM-1-(+); Ki67-(+) cardiomyocyte nuclei. N. S., not significant. *, P-value < 0.05. **, P-value < 0.01. Scale bar represents 50 μm.

To determine if increased mononucleated cardiomyocytes in *TH-Cre; TrkA^f/f^* mice correlated with increased cardiomyocyte proliferation, we analyzed 5 μm-thick cyrosections from mutant and control hearts (Fig. 1C-D). To unambiguously identify cardiomyocyte nuclei, we stained tissue sections with antibodies against pericentriolar material 1 (PCM-1), which adopts a perinuclear localization specifically in cardiomyocytes.^46^ Additionally, sections were stained with DAPI to visualize nuclei and antibodies against Ki67, a proliferative marker, to locate nuclei within proliferating cells. Quantification of PCM-1-(+); Ki67-(+) proliferative cardiomyocytes at P14 revealed that while *TrkA^f/f^* control mice exhibited low levels of basal cardiomyocyte proliferation (0.5%, n = 3), cardiomyocyte proliferation was significantly increased in *TH-Cre; TrkA^f/f^* mice (1.29%, n = 3). Taken together, these results demonstrate that ablation of sympathetic nerves increases the percentage of mononucleated cardiomyocytes with a corresponding increase in cardiomyocyte proliferation.

### β-adrenergic receptor inhibition increases cardiomyocyte mononucleation and proliferation

Through the release of potent neurotransmitters such as norepinephrine (NE), sympathetic neurons activate β-adrenergic receptors (βARs) expressed in innervated tissues.^47^ To test if βAR-activation affects cardiomyocyte nucleation and proliferation during perinatal development, we injected mice once-a-day from P0 to P14 with propranolol (40 μg per g bodyweight), a non-selective β-blocker.^48^ We harvested hearts from propranolol-injected mice at P14 and dissociated cardiomyocytes into isolated single cells (Fig. 2A-B). Analysis and quantification of cardiomyocyte nucleation revealed that βAR inhibition during the first two weeks after birth increased the percentage of mononucleated cardiomyocytes (22.1%, n = 6) compared to that observed in saline-injected controls (11.6%, n = 7). Additionally, when we analyzed the hearts of one-year-old mutant mice lacking function of all three βAR isoforms (βAR TKO) we again observed an increased percentage of mononucleated cardiomyocytes in these mutants (17.6%, n = 3) compared to that observed in age-matched wild-type mice (5.19%, n = 3) (Fig. 2C-D).

**Figure 2.**
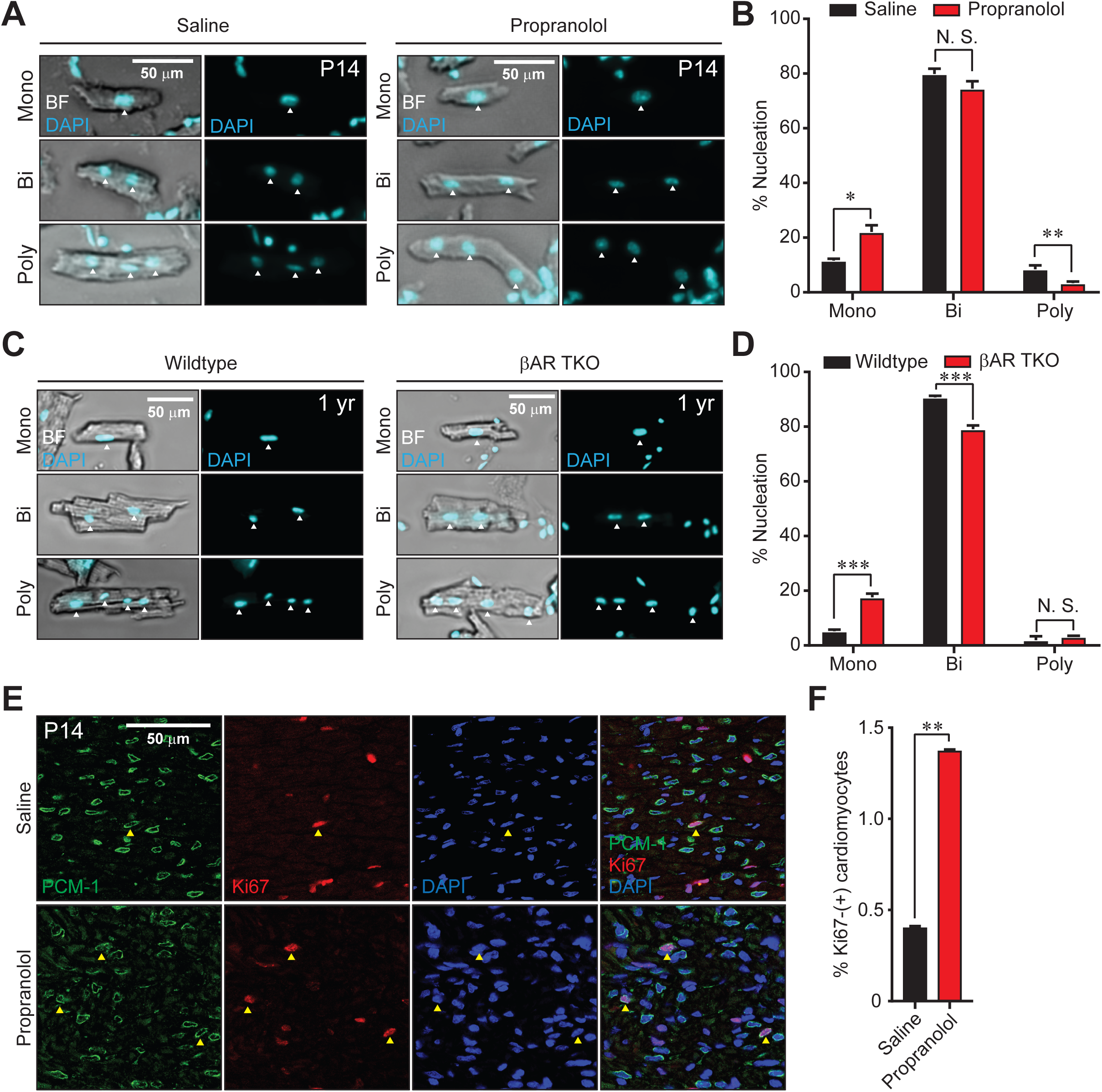
Inhibition of βARs increases cardiomyocyte mononucleation and proliferation *in vivo*. **A)** Representative micrographs and **B)** quantification of mononucleated (Mono), binucleated (Bi), and polynucleated (Poly) cardiomyocytes dissociated from P14 propranolol- (n = 6) and saline-injected (n= 7) mice (once per day, P0 to P14). **C)** Representative micrographs and **D)** quantification of Mono, Bi, and Poly cardiomyocytes dissociated from one-year-old βAR TKO (n = 3) and wild-type control (n = 3) mice. White arrowheads indicate cardiomyocyte-specific nuclei. **E)** Representative micrographs and **F)** quantification of 5 μm-thick sections from P14 hearts harvested from propranolol- or saline-injected mice stained with PCM-1 (cardiomyocyte nuclei), Ki67 (proliferative cells), and DAPI (all nuclei). Yellow arrowheads indicate PCM-1-(+); Ki67-(+) cardiomyocyte nuclei. N. S., not significant. *, P-value < 0.05. **, P-value < 0.01, ***, P-value < 0.001. Scale bar represents 50 μm.

To assess if elevated mononucleated cardiomyocytes in animals with reduced βAR function correlated to increased cardiomyocyte proliferation, we harvested propranolol- and saline-injected hearts at P14, generated 5 μm-thick cryosections, and stained these samples with antibodies against PCM-1 and Ki67. Quantification of PCM-1-(+); Ki67-(+) cells indicated that inhibition of βARs during the perinatal window with daily propranolol injection increased the percentage of proliferative cardiomyocytes (1.38%, n = 3) compared to saline-injected controls (0.41%, n = 3). These results suggest that inhibition of βARs specifically during first two weeks after birth results in increased numbers of mononucleated cardiomyocytes with increased proliferative activity *in vivo*, corroborating the phenotypes observed in hearts lacking sympathetic nerve innervation.

### Activation of βARs suppress cardiomyocyte proliferation in vitro

Though our *in vivo* experiments suggested a role for sympathetic nerve activity in cardiomyocyte cell-cycle control, it is possible that these effects are mediated through indirect mechanisms as the heart consists of other cell types including fibroblasts and endothelial cells.^49^ To more directly test the effects of bAR signaling on cardiomyocytes themselves, we isolated and cultured cardiomyocytes from P1-P3 neonatal mice. By analyzing incorporation of the thymidine nucleoside analog 5-ethynyl-2’-deoxyuridine (EdU) as a marker for proliferative activity combined with antibody staining of PCM-1 and the cardiomyocyte-specific marker cardiac troponin T (cTnT), we characterized the basal proliferation rates of cardiomyocytes in our *in vitro* culture system (Fig. 3A). From six independent cardiomyocyte culture experiments, we determined that on average ∼1.5% of cardiomyocytes are EdU-(+) after 2 days of incubation with EdU; however, we also noticed that baseline proliferation among independent experiments ranged greatly (0.76% to 2.26%) (Fig. 3B). To take this variation into account, we normalized all experimental conditions against the control baseline proliferation for that specific experiment.

**Figure 3.**
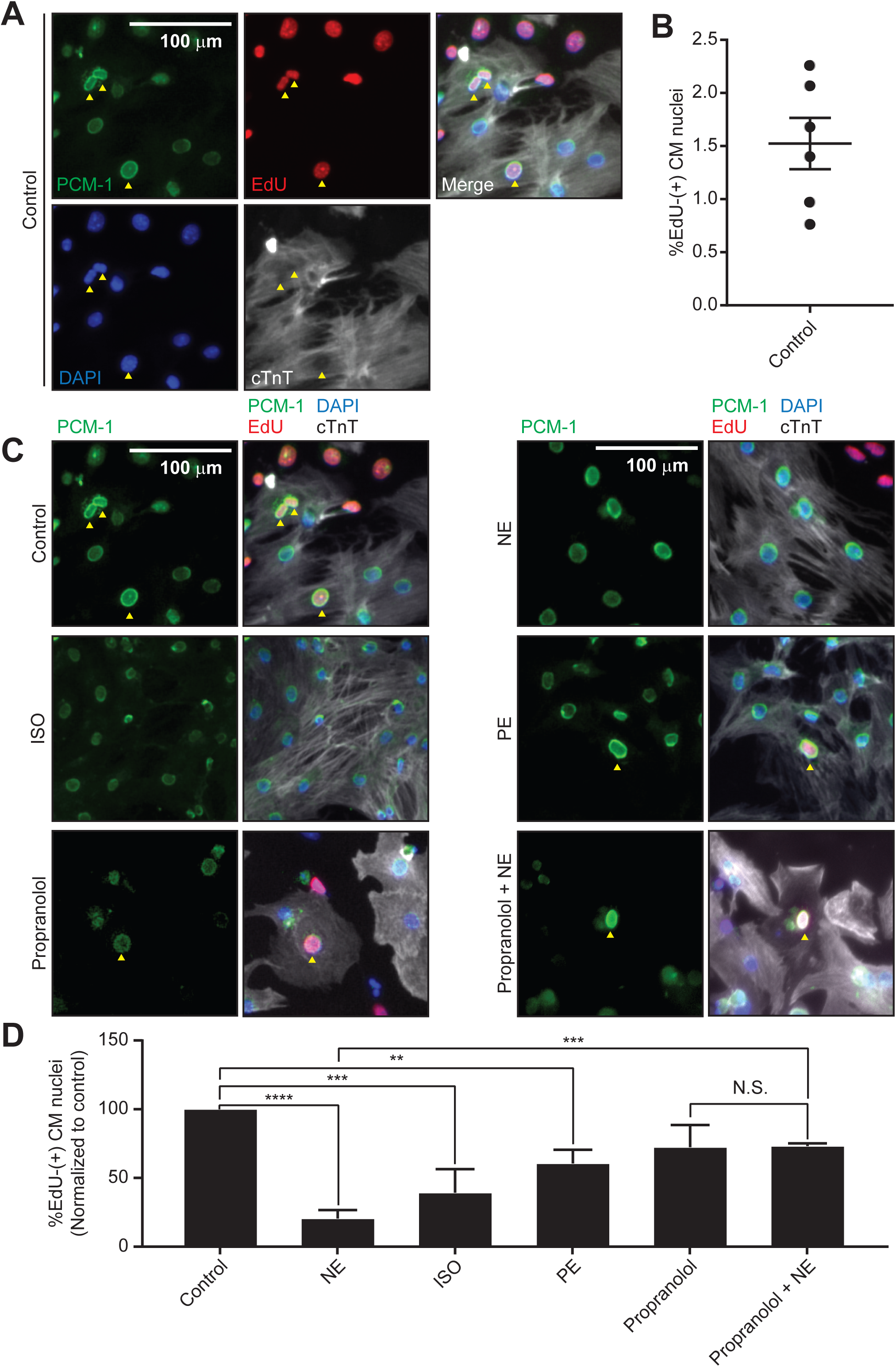
Activation of βAR inhibits cardiomyocyte proliferation *in vitro*. **A)** Representative micrographs of neonatal cardiomyocyte cultures derived from P1 to P3 mice labeled with EdU (proliferative cells), DAPI (nuclei), and antibodies against PCM-1 (cardiomyocyte nuclei) and cTnT (cardiomyocyte). **B)** Basal proliferation observed in six independent cardiomyocyte culture experiments. **C)** Representative micrographs and **D)** quantification of cardiomyocyte proliferation observed after incubation of cultures with 10 μM norepinephrine (NE), 10 μM isoproterenol (ISO), 100 μM phenylephrine (PE), and 10 μM propranolol. Yellow arrowheads indicate PCM-1-(+); cTnT-(+); EdU-(+) proliferating cardiomyocytes. Results from three biological replicates are shown. N. S., not significant. *, P-value < 0.05. **, P-value < 0.01, ***, P-value < 0.001, **** P-value < 0.0001. Scale bar represents 100 μm.

We tested a panel of chemical modulators of both β- and α-adrenergic receptors in our *in vitro* proliferation assay (Fig. 3C-D). Activation of both β- and α-adrenergic receptors with norepinephrine (NE) resulted in strong suppression of cardiomyocyte proliferation. βAR-specific activation with isoproterenol (ISO) also caused significant inhibition of cardiomyocyte cell-cycle activity, supporting the role of βAR in cardiomyocyte proliferative control. Treatment with phenylephrine (PE), an α-adrenergic receptor (αAR)-specific agonist, also resulted in mild suppression of cardiomyocyte cell-cycle activity possibly suggesting at least some role for αARs in controlling cardiomyocyte proliferation. Importantly, treatment with the βAR inhibitor propranolol in combination with NE prevented NE-dependent inhibition of cardiomyocyte proliferation, confirming that NE exerts its effects on cardiomyocyte proliferation through βARs. In combination, our *in vitro* results are in agreement with our *in vivo* findings illustrating a role for sympathetic nerve activity through regulation of βARs in cardiomyocyte cell-cycle control.

### βARs function through PKA to control cardiomyocyte cell-cycle activity

Canonical βAR activation is believed to increase intracellular cyclic AMP levels leading to activation of cyclic AMP-dependent protein kinase (PKA) to mediate downstream cellular effects.^50^ We hypothesized that βAR-dependent regulation of cardiomyocyte proliferation could also depend on PKA activity. Using our *in vitro* proliferation assay, we treated cultured cardiomyocytes with NE alone and in combination with chemical modulators of PKA signaling, forskolin and KT5720 (Fig. 4A-B). Treatment of cardiomyocytes with the PKA activator forskolin phenocopied the effects of NE treatment leading to robust inhibition of cardiomyocyte proliferation. While addition of the PKA inhibitor KT5720 did not have a significant effect on baseline cardiomyocyte proliferation, treatment of both KT5720 and NE simultaneously reduced the efficacy of NE to inhibit cardiomyocyte proliferation. Additionally, treatment of NE with forskolin did not lead to enhanced cell-cycle suppression beyond that observed with NE treatment alone, suggesting that NE may function predominantly through a PKA-dependent mechanism to regulate cardiomyocyte cell-cycle activity.

**Figure 4.**
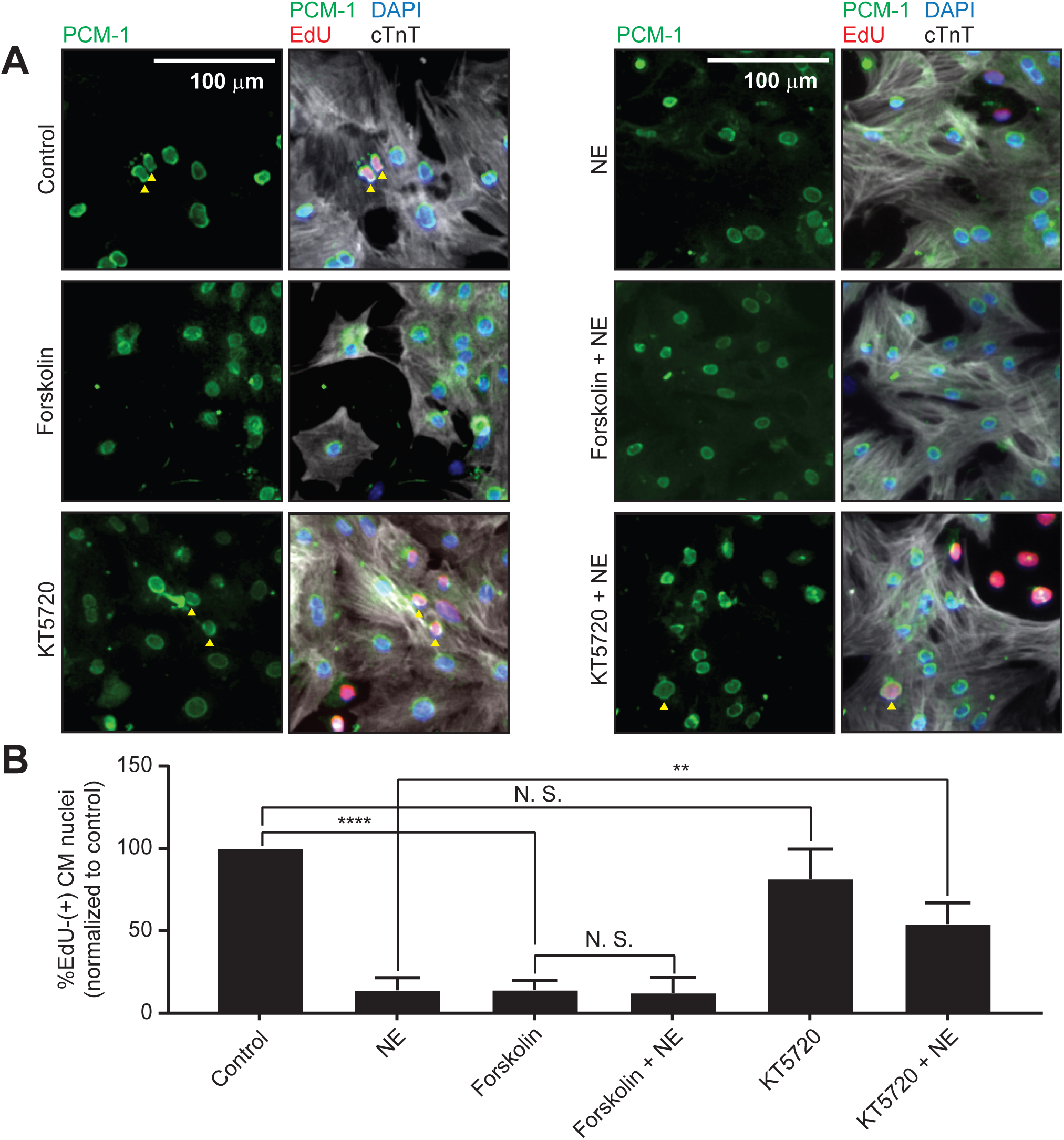
βAR-dependent control of cardiomyocyte proliferation functions requires PKA. **A)** Representative micrographs and **B)** quantification of cardiomyocyte proliferation observed after incubation of cultures with 10 μM norepinephrine (NE), 1 μM forsokolin, and 1 μM KT5720. Cultures are labeled with EdU (proliferative cells), DAPI (nuclei), and antibodies against PCM-1 (cardiomyocyte nuclei) and cTnT (cardiomyocyte). Yellow arrowheads indicate PCM-1-(+); cTnT-(+); EdU-(+) proliferating cardiomyocytes. Results from three biological replicates are shown. N. S., not significant. **, P-value < 0.01, **** P-value < 0.0001. Scale bar represents 100 μm.

## DISCUSSION

Cumulatively, our results support a model where sympathetic nerve activity during neonatal development promotes cardiomyocyte cell-cycle arrest and binucleation (Fig. 5). Using a genetic system to ablate sympathetic neurons, we demonstrate that mice lacking sympathetic innervation possess elevated mononucleated cardiomyocyte populations with increased proliferative activity two weeks after birth. Furthermore, by examining mutant mice lacking function of all three βAR isoforms, we observed a similar increase in cardiomyocyte mononucleation that persisted until at least one year of age, suggesting that sympathetic nerves may act through βARs to mediate cardiomyocyte cell-cycle control *in vivo*. Pharmacological inhibition of βARs with propranolol specifically within the first two weeks after birth phenocopied the increase in cardiomyocyte mononucleation and proliferation observed in animals depleted of sympathetic neurons. Therefore, despite cardiac sympathetic nerve innervation at E13.5,^38^ it is likely that nerve-dependent activation of βARs during the first two-weeks after birth is responsible for limiting cardiomyocyte proliferative capacity. Our *in vitro* cardiomyocyte culture experiments corrborate our *in vivo* findings demonstrating that activation of βARs inhibits neonatal cardiomyocyte proliferation through a PKA-dependent mechanism.

**Figure 5.**
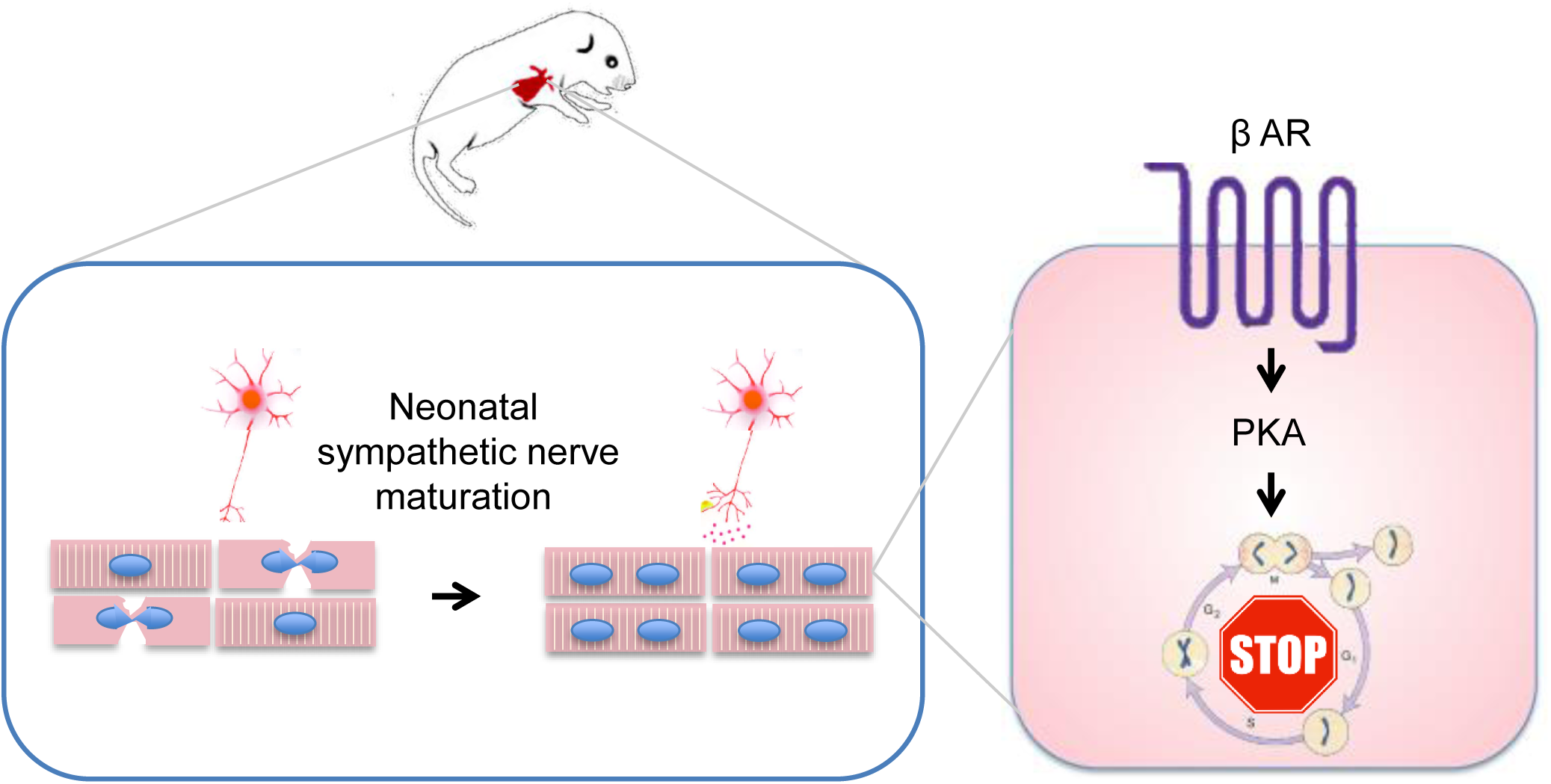
A model for sympathetic nerve-dependent cardiomyocyte cell-cycle control. Sympathetic nerves promote cardiomyocyte binucleation during the neonatal window. Release of neurotransmitters activates βAR receptors expressed in cardiomyocytes which functions through PKA to inhibit cell-cycle activity.

Activation of βARs is known to regulate the rate and force of cardiomyocyte contractility.^37^ During adult zebrafish heart regeneration, cardiomyocyte dedifferentiation and sarcomeric disassembly have been observed suggesting that inactivation of the contractile apparatus may be necessary for cell division.^51^ It is possible that the increase in cardiac sympathetic tone during the first weeks after birth may stabilize the contractile machinery preventing cell proliferation. However while treatment of neonatal cardiomyocytes with the αAR-specific agonist phenylephrine dramatically increased the rate of cardiomyocyte contraction *in vitro* (data not shown), only mild reduction of cardiomyocyte proliferation was observed in our assays (Fig 3C-D). Therefore, βARs could potentially regulate cardiomyocyte cell-cycle independently from contractility.

It was previously reported that treatment of neonatal rats with propranolol inhibited cardiomyocyte proliferation through the P70 S6K pathway.^52^. It is important to note that while we focused on the effect of βAR inhibition over the course of two-weeks, Tseng et al focused on the effects of βAR inhibition over the time scale of minutes to hours. Additionally, to increase the accuracy of our quantifications, we stained our tissues and cells with PCM-1, a recently validated marker for cardiomyocyte nuclei.^46^ Nevertheless, it remains possible that inhibiting βAR signaling could have differing effects on cardiomyocyte proliferation depending on the timescale of inhibition.

Two groups have recently highlighted the importance of nerves in neonatal mouse heart regeneration.^53,54^ Mahmoud et al demonstrated the importance of parasympathetic nerves in the regulation of cardiomyocyte proliferation and regeneration in both zebrafish and neonatal mice. Inhibition of cholinergic transmission by treatment with the non-selective muscarinic receptor antagonist atropine decreased cardiomyocyte proliferation.^53^ Importantly, treatment with propranolol did not inhibit cardiomyocyte proliferation during regeneration. Additionally, surgical ablation of the left vagus nerve inhibited proliferation and regeneration. Combined these results strongly support the importance of parasympathetic inputs during cardiac regeneration. In a recent study, White et al reported the requirement of sympathetic nerve reinnervation during neonatal mouse heart regeneration.^54^ Chemical sympathectomy was performed by injecting neonatal mice with 6-hydroxydopamine hydrobromide (6-OHDA), a neurotoxin that induces apoptosis in catecholaminergic neurons, which inhibited cardiac regeneration and promoted fibrosis. While robust denervation was observed in the heart, it is possible that the non-specificity of 6-OHDA treatment could induce apoptosis of other catecholaminergic terminals, such as in the central nervous system, that could affect heart regeneration. Therefore, from this data alone, it is difficult to assess the importance of sympathetic nerves in the context of cardiac regeneration.

The physiological events triggering cardiomyocyte cell-cycle arrest and loss of regenerative potential in mammals has remained enigmatic. In this study, we report that sympathetic nerve activity through βARs is one the triggers promoting cardiomyocyte cell-cycle exit and loss of proliferative capacity. While we observe increased cardiomyocyte mononucleation and proliferation in animals with perturbed βAR signaling, we also notice that the majority of cardiomyocytes (∼80%) still underwent binucleation by P14. This implies that factors other than the sympathetic nervous system must also act to promote cardiomyocyte cell-cycle arrest. Identification of these new signals will be important as it is likely that modulation of multiple pathways in combination will be necessary to unlock regenerative potential in adult mammals, which will provide us with new therapeutic strategies in regenerative medicine.

## ACKNOWLEDGEMENTS

This work was supported by UCSF-IRACDA postdoctoral fellowship (A. P.), and NIH Pathway to Independence Award, Edward Mallinckrodt Jr. Foundation, March of Dimes Basil O’Conner Scholar Award, American Heart Association, American Federation for Aging Research, Life Sciences Research Foundation, Program for Breakthrough Biomedical Research, UCSF Eli and Edythe Broad Center of Regeneration Medicine and Stem Cell Research, Resource Allocation Program, and Cardiovascular Research Institute (G. H.).

## COMPETING FINANCIAL INTERESTS

The authors declare no competing financial interests.

## REFERENCES

1 Xin, M., Olson, E. N. & Bassel-Duby, R. Mending broken hearts: cardiac development as a basis for adult heart regeneration and repair. Nat. Rev. Mol. Cell Biol. 14, 529–541 (2013).

2 Oberpriller, J. O. & Oberpriller, J. C. Response of the adult newt ventricle to injury. J. Exp. Zool. 187, 249–259 (1974).

3 Bader, D. & Oberpriller, J. O. Repair and reorganization of minced cardiac muscle in the adult newt (Notophthalmus viridescens). J. Morphol. 155, 349–357 (1978).

4 Bader, D. & Oberpriller, J. Autoradiographic and electron microscopic studies of minced cardiac muscle regeneration in the adult newt, Notophthalmus viridescens. J. Exp. Zool. 208, 177–193 (1979).

5 Witman, N., Murtuza, B., Davis, B., Arner, A. & Morrison, J. I. Recapitulation of developmental cardiogenesis governs the morphological and functional regeneration of adult newt hearts following injury. Dev. Biol. 354, 67–76 (2011).

6 Claycomb, W. C. Cardiac-muscle hypertrophy. Differentiation and growth of the heart cell during development. Biochem. J. 168, 599–601 (1977).

7 Poss, K. D., Wilson, L. G. & Keating, M. T. Heart Regeneration in Zebrafish. Science 298, 2188–2190 (2002).

8 Choi, W.-Y. & Poss, K. D. Cardiac Regeneration. Curr. Top. Dev. Biol. 100, 319–344 (2012).

9 Porrello, E. R. et al. Transient regenerative potential of the neonatal mouse heart. Science 331, 1078–1080 (2011).

10 Haubner, B. J. et al. Functional Recovery of a Human Neonatal Heart After Severe Myocardial Infarction. Circ. Res. 118, 216–221 (2016).

11 Porrello, E. R. et al. Regulation of neonatal and adult mammalian heart regeneration by the miR-15 family. Proc. Natl. Acad. Sci. U. S. A. 110, 187–192 (2013).

12 Senyo, S. E. et al. Mammalian heart renewal by pre-existing cardiomyocytes. Nature 493, 433–436 (2013).

13 Huang, G. N. et al. C/EBP transcription factors mediate epicardial activation during heart development and injury. Science 338, 1599–1603 (2012).

14 Muneoka, K., Allan, C. H., Yang, X., Lee, J. & Han, M. Mammalian regeneration and regenerative medicine. Birth Defects Res. Part C Embryo Today Rev. 84, 265–280 (2008).

15 Wang, J. M., Irwin, R. W., Liu, L., Chen, S. & Brinton, R. D. Regeneration in a degenerating brain: potential of allopregnanolone as a neuroregenerative agent. Curr. Alzheimer Res. 4, 510–517 (2007).

16 Walmsley, G. G. et al. Murine Dermal Fibroblast Isolation by FACS. J. Vis. Exp. JoVE (2016). doi:10.3791/53430

17 Wilgus, T. A. Regenerative healing in fetal skin: a review of the literature. Ostomy. Wound Manage. 53, 16–31; quiz 32-33 (2007).

18 Wulff, B. C. et al. Mast cells contribute to scar formation during fetal wound healing. J. Invest. Dermatol. 132, 458–465 (2012).

19 Kikuchi, K. et al. Primary contribution to zebrafish heart regeneration by gata4(+) cardiomyocytes. Nature 464, 601–605 (2010).

20 Jopling, C. et al. Zebrafish heart regeneration occurs by cardiomyocyte dedifferentiation and proliferation. Nature 464, 606–609 (2010).

21 Wills, A. A., Holdway, J. E., Major, R. J. & Poss, K. D. Regulated addition of new myocardial and epicardial cells fosters homeostatic cardiac growth and maintenance in adult zebrafish. Dev. Camb. Engl. 135, 183–192 (2008).

22 Bettencourt-Dias, M., Mittnacht, S. & Brockes, J. P. Heterogeneous proliferative potential in regenerative adult newt cardiomyocytes. J. Cell Sci. 116, 4001–4009 (2003).

23 Kikuchi, K. et al. Retinoic acid production by endocardium and epicardium is an injury response essential for zebrafish heart regeneration. Dev. Cell 20, 397–404 (2011).

24 Xin, M. et al. Regulation of insulin-like growth factor signaling by Yap governs cardiomyocyte proliferation and embryonic heart size. Sci. Signal. 4, ra70 (2011).

25 von Gise, A. et al. YAP1, the nuclear target of Hippo signaling, stimulates heart growth through cardiomyocyte proliferation but not hypertrophy. Proc. Natl. Acad. Sci. U. S. A. 109, 2394–2399 (2012).

26 Heallen, T. et al. Hippo signaling impedes adult heart regeneration. Dev. Camb. Engl. 140, 4683–4690 (2013).

27 Xin, M. et al. Hippo pathway effector Yap promotes cardiac regeneration. Proc. Natl. Acad. Sci. U. S. A. 110, 13839–13844 (2013).

28 Lin, Z. et al. Cardiac-specific YAP activation improves cardiac function and survival in an experimental murine MI model. Circ. Res. 115, 354–363 (2014).

29 Xiang, F.-L., Guo, M. & Yutzey, K. E. Overexpression of Tbx20 in Adult Cardiomyocytes Promotes Proliferation and Improves Cardiac Function After Myocardial Infarction. Circulation 133, 1081–1092 (2016).

30 Chakraborty, S., Sengupta, A. & Yutzey, K. E. Tbx20 promotes cardiomyocyte proliferation and persistence of fetal characteristics in adult mouse hearts. J. Mol. Cell. Cardiol. 62, 203–213 (2013).

31 Chakraborty, S. & Yutzey, K. E. Tbx20 regulation of cardiac cell proliferation and lineage specialization during embryonic and fetal development in vivo. Dev. Biol. 363, 234–246 (2012).

32 Stennard, F. A. et al. Murine T-box transcription factor Tbx20 acts as a repressor during heart development, and is essential for adult heart integrity, function and adaptation. Dev. Camb. Engl. 132, 2451–2462 (2005).

33 Singh, M. K. et al. Tbx20 is essential for cardiac chamber differentiation and repression of Tbx2. Dev. Camb. Engl. 132, 2697–2707 (2005).

34 Takeuchi, J. K. et al. Tbx20 dose-dependently regulates transcription factor networks required for mouse heart and motoneuron development. Dev. Camb. Engl. 132, 2463–2474 (2005).

35 Mahmoud, A. I. et al. Meis1 regulates postnatal cardiomyocyte cell cycle arrest. Nature 497, 249–253 (2013).

36 Puente, B. N. et al. The oxygen-rich postnatal environment induces cardiomyocyte cell-cycle arrest through DNA damage response. Cell 157, 565–579 (2014).

37 Triposkiadis, F. et al. The Sympathetic Nervous System in Heart Failure: Physiology, Pathophysiology, and Clinical Implications. J. Am. Coll. Cardiol. 54, 1747–1762 (2009).

38 Manousiouthakis, E., Mendez, M., Garner, M. C., Exertier, P. & Makita, T. Venous endothelin guides sympathetic innervation of the developing mouse heart. Nat. Commun. 5, 3918 (2014).

39 Bartolomé, J., Lau, C. & Slotkin, T. A. Ornithine decarboxylase in developing rat heart and brain: role of sympathetic development for responses to autonomic stimulants and the effects of reserpine on maturation. J. Pharmacol. Exp. Ther. 202, 510–518 (1977).

40 Soonpaa, M. H., Kim, K. K., Pajak, L., Franklin, M. & Field, L. J. Cardiomyocyte DNA synthesis and binucleation during murine development. Am. J. Physiol. 271, H2183–2189 (1996).

41 Borden, P., Houtz, J., Leach, S. D. & Kuruvilla, R. Sympathetic innervation during development is necessary for pancreatic islet architecture and functional maturation. Cell Rep. 4, 287–301 (2013).

42 Bachman, E. S. et al. betaAR signaling required for diet-induced thermogenesis and obesity resistance. Science 297, 843–845 (2002).

43 Judd, J., Lovas, J. & Huang, G. N. Isolation, Culture and Transduction of Adult Mouse Cardiomyocytes. J. Vis. Exp. JoVE (2016). doi:10.3791/54012

44 Sander, V., Suñe, G., Jopling, C., Morera, C. & Izpisua Belmonte, J. C. Isolation and in vitro culture of primary cardiomyocytes from adult zebrafish hearts. Nat. Protoc. 8, 800–809 (2013).

45 Majdan, M., Walsh, G. S., Aloyz, R. & Miller, F. D. TrkA mediates developmental sympathetic neuron survival in vivo by silencing an ongoing p75NTR-mediated death signal. J. Cell Biol. 155, 1275–1286 (2001).

46 Alkass, K. et al. No Evidence for Cardiomyocyte Number Expansion in Preadolescent Mice. Cell 163, 1026–1036 (2015).

47 Grassi, G., Mark, A. & Esler, M. The Sympathetic Nervous System Alterations in Human Hypertension. Circ. Res. 116, 976–990 (2015).

48 Bn, P. Propranolol in the treatment of angina: a review. Postgrad. Med. J. 52 Suppl 4, 35–41 (1976).

49 Camelliti, P., Borg, T. K. & Kohl, P. Structural and functional characterisation of cardiac fibroblasts. Cardiovasc. Res. 65, 40–51 (2005).

50 Hausdorff, W. P., Caron, M. G. & Lefkowitz, R. J. Turning off the signal: desensitization of beta-adrenergic receptor function. FASEB J. 4, 2881–2889 (1990).

51 Kikuchi, K. Advances in understanding the mechanism of zebrafish heart regeneration. Stem Cell Res. 13, 542–555 (2014).

52 Tseng, Y.-T. et al. β-Adrenergic receptors (βAR) regulate cardiomyocyte proliferation during early postnatal life. FASEB J. 15, 1921–1926 (2001).

53 Mahmoud, A. I. et al. Nerves Regulate Cardiomyocyte Proliferation and Heart Regeneration. Dev. Cell 34, 387–399 (2015).

54 White, I. A., Gordon, J., Balkan, W. & Hare, J. M. Sympathetic Reinnervation is Required for Mammalian Cardiac Regeneration. Circ. Res. CIRCRESAHA.115.307465 (2015). doi:10.1161/CIRCRESAHA.115.307465

